# Reactivated past decisions repel early sensory processing and attract late decision-making

**DOI:** 10.1101/2024.02.26.582221

**Authors:** Minghao Luo, Huihui Zhang, Huan Luo

## Abstract

Automatic shaping of perception by past experiences is common in many cognitive functions, reflecting the exploitation of temporal regularities in environments. A striking example is serial dependence, i.e., current perception is biased by previous trials. However, the neural implementation of its operational circle in human brains remains unclear. In two experiments with Electroencephalography (EEG) / Magnetoencephalography (MEG) recordings and delayed-response tasks, we demonstrate a two-stage ’repulsive-then-attractive’ past-present interaction mechanism underlying serial dependence. First, past-trial reports serve as a prior to be reactivated during both encoding and decision-making. Crucially, past reactivation interacts with current information processing in a two-stage manner: repelling and attracting the present during encoding and decision-making, and arising in the sensory cortex and prefrontal cortex, respectively. Finally, while the early stage occurs automatically, the late stage is modulated by task and predicts bias behavior. Our findings might also illustrate general mechanisms of past-present influences in neural operations.

## Introduction

Although the outside world is changing constantly, natural stimuli maintain regularities over time^1,2^. Accordingly, perception is not solely determined by current sensory inputs, but largely shaped by past experiences, reflecting the exploitation of temporal regularities in the environment^3^. One striking example is serial dependence, that is, perception tends to be involuntarily biased towards stimuli in previous trials^4–8^. Serial dependence has been robustly observed in a wide range of visual features, from simple ones like orientation^4,9–12^, numerosity^13^, and position^14,15^ to abstract ones such as facial attractiveness^16,17^ and other modalities^18,19^. Despite its detrimental effects on perceptual fidelity, serial dependence is thought to reflect an optimizing strategy of the brain to increase perceptual stability and efficiency by taking advantage of inherent temporal correlations in a world that is presumably stable over a short time scale^20,21^.

Two major theories have been proposed to account for serial dependence. The “continuity field” view posits that serial dependence occurs at the perceptual level whereby similar stimuli of spatiotemporal adjacency are automatically integrated, leading to the typically observed attractive serial bias^4^. In contrast, the post-perceptual view^11,22^ postulates a two-stage process consisting of perceptual-level efficient coding and post-perceptual Bayesian inference, associated with repulsive and attractive serial bias, respectively.

Although large amounts of behavioral experiments have been performed to address the issue^9–11,23,24^, neural evidence is certainly crucial to reconciling the arguments, which remain mixed. For example, neural activities in primary visual cortex have been found to be shifted toward previous trials, advocating the perceptual “continuity field” view^25^, yet challenged by other studies revealing repulsive neural bias in the visual cortex despite attractive serial bias in behavior^26,27^. Moreover, animal electrophysiological studies and human Transcranial Magnetic Stimulation (TMS) studies demonstrate the involvement of higher-level regions, such as the prefrontal cortex (PFC) and posterior parietal cortex (PPC), in attractive serial dependence^28–30^, thus commensurate with the post-perceptual view. Overall, it remains unclear how serial dependence dynamically emerges along various stages of processing, i.e., from early sensory regions to higher-level regions, and ultimately contributes to bias behavior.

In addition to the theoretical frameworks focusing on the stages of its occurrence, serial dependence could also be understood as neural interactions between memory reactivations and current information processing, given its automatic influence across independent trials. PFC recordings on monkeys show that past-trial information is transiently reactivated just before trial onset and further correlates with serial bias behavior^28^. Our recent study^19^ as well as previous works^31–34^ demonstrate that past-trial information, indeed retained in activity-silent states during the inter-trial interval, is triggered by the corresponding event within the current trial, and the co-occurrence of past and present contributes to serial bias. Most importantly, the past-to-present neural influence exhibits a feature-specific direction commensurate with the corresponding bias behavior, i.e., repulsive for sensory and motor features, and attractive for choices^19^. Serial bias is also posited to depend on neural representational similarities between sensory and memory information^35^, such as aligned and flipped relationships resulting in attractive and repulsive serial bias^26,33^. Therefore, elucidating past-present interactions and the respective shifting directions in multiple brain regions is vital to understanding neural implementations of serial dependence.

The present study aims to unravel when and where serial dependence occurs by systematically examining past-trial reactivation and its interaction with present information processing in the whole brain. Crucially, we developed a delayed-response task to explicitly separate perception and decision-making stages so that their intertwined roles could be disambiguated. Behaviorally, participants showed attractive serial bias towards previously reported locations in continuous 2-D spatial perception. EEG/MEG recordings in two experiments demonstrate that the past-trial report information is reactivated and interacts with present information processing during both encoding and decision-making stages but in distinct ways, i.e., repulsive and automatic during encoding, while attractive, modulated by task relevance and correlated with behavior during decision-making, originated in the visual cortex and prefrontal cortex, respectively. Taken together, serial dependence arises from a two-stage “repulsive-followed-by-attractive” past-present interaction process, which might contribute to a wide range of themes beyond perception.

## Results

### Attractive serial dependence in 2-D continuous spatial perception

In Experiment 1, 29 human participants performed a delayed-response spatial location reproduction task with their 64-channel EEG activities recorded (Fig. 1A). Specifically, a red dot was presented within a 2-D round area (Encoding stage) followed by a square noise mask, and participants reproduced the target location using a mouse starting at a random location (Decision-making stage). A multiple linear regression analysis (Fig. 1B) revealed that the 2-D spatial reproduction performance (“Current Report”) was modulated by stimulus location (“Current Stimulus”; *t*(28) = 84.77, p < 0.001), starting location of the mouse (“Current Start”; *t*(28) = 4.56, p < 0.001), and most importantly, the reported location in the preceding trial (“Previous Report”; *t*(28) = 2.78, p = 0.0096), indicating an attractive serial bias in behavior, i.e., the reported location was biased towards that of the previous trial.

**Figure 1.**
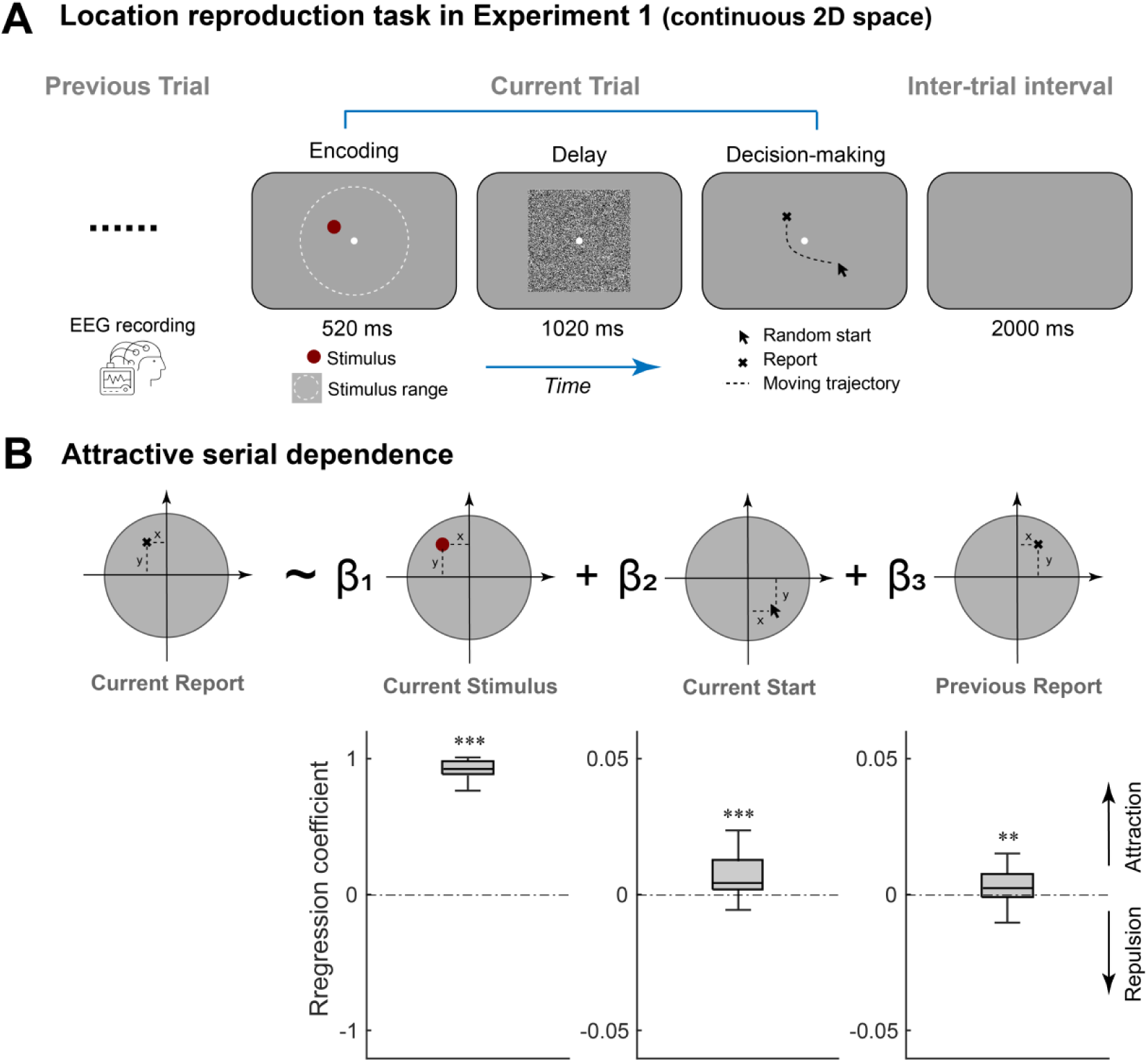
Attractive serial bias behavior in 2-D spatial perception task (Experiment 1, EEG). **A**. On each trial, a red dot was randomly presented within a 2-D round area (Encoding period), followed by a square noise mask (Delay period). During Decision-making period, participants moved the mouse starting at a random location (Random start) to reproduce the red dot location, with their EEG activities recorded. **B**. Upper: a trial-wise linear regression model accounting for the reproduced location based on 3 factors: location of current target (current stimulus), starting location of mouse cursor (Current Start), reproduced location in previous trial (Previous Report). Lower: box plots of the regression coefficients (Left to right: Current stimulus, Current Start, Previous Report). Edges from bottom to top indicate 25^th^, 50^th^ and 75^th^ percentiles, respectively. (**: p < 0.01, and ***: p < 0.001).

### Two-stage reactivations of previous-trial information

As serial dependence engages automatic past-to-present modulation, past information should be carried over and coexist with the current stimulus in the present trial. We therefore first sought to decode the stimulus location of preceding trial (N-1) and current trial (N) from the current-trial brain activities, by conducting a trial-wise representational similarity analysis (RSA) at each time point and on each participant (Fig. 2A). Specifically, the dissimilarity (distance) of target locations within 2-D continuous space was calculated for each trial pair, based on which the representational dissimilarity matrix (RDM) was constructed (Location RDM). The neural dissimilarity matrix (Neural RDM) based on multivariate EEG patterns was then regressed by Location RDM, resulting in representational strength (Fig. 2A, right). Notably, the RSA analysis was performed for current-trial and previous-trial location RDMs separately, resulting in the corresponding neural decoding time courses (Fig. 2B, right).

**Figure 2.**
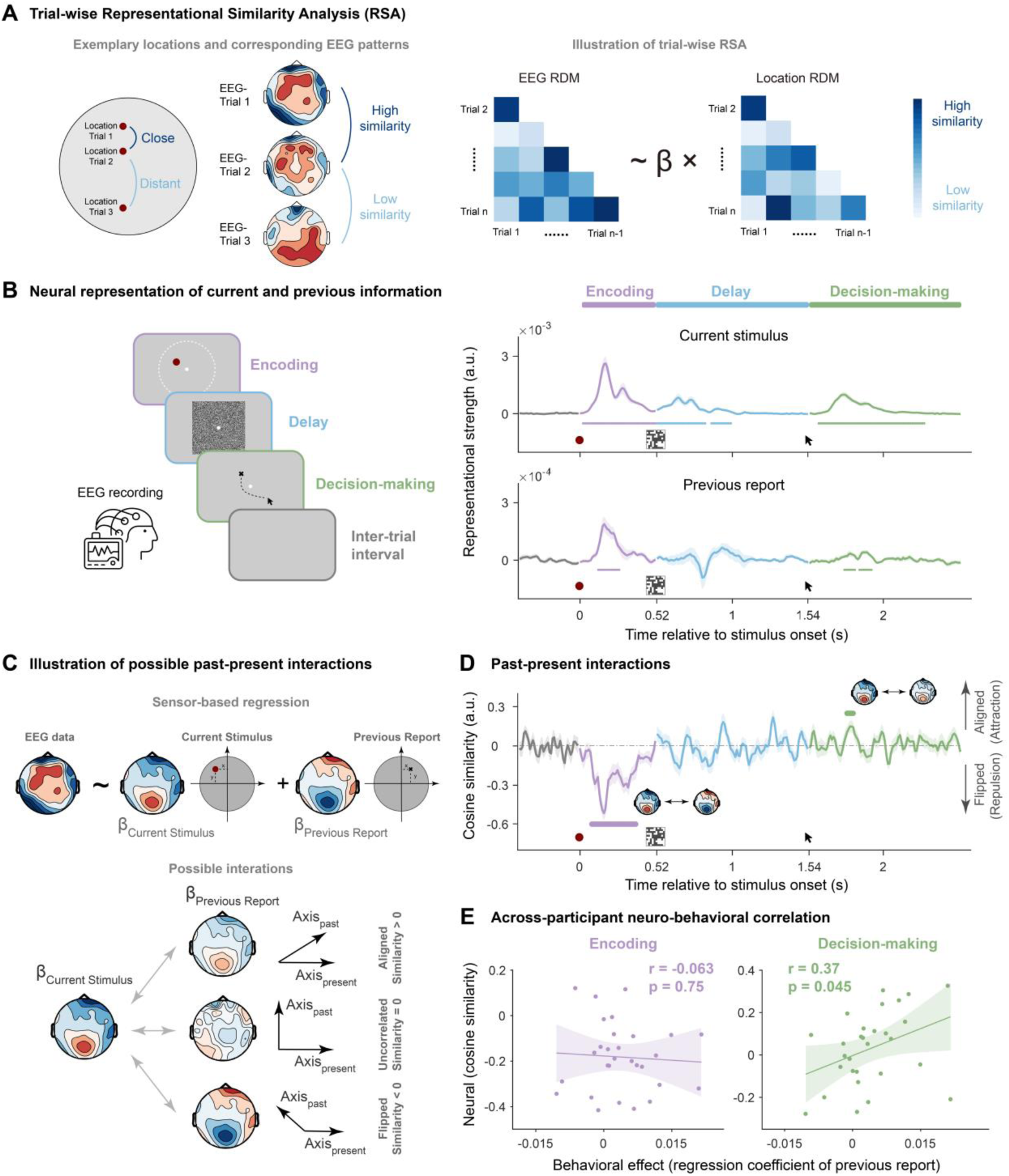
Neural decoding of current-trial and past-trial information, past-present neural interaction and its correlates to behavior (Experiment 1). **A.** Schematic illustration of trial-wise RSA (representational similarity analysis). Left: For every pair of trials, closer distance in dot location (dark blue) is hypothesized to have more similar multivariate EEG pattern than that with far spatial distance (light blue), yielding the location RDM (right panel). Right: EEG RDM is calculated based on the multivariate EEG similarity for each pair of trials. Regression of EEG RDM into location RDM yields the regression coefficient (β) denoting the neural representation strength of location information. **B**. Left: Each trial consists of three stages: Encoding (purple), Delay (blue), and Decision-making (green). Right: Time-resolved grand averaged decoding of current-trial stimulus (upper) and past-trial reported location (lower). Note that the three stages – Encoding (purple), Delay (blue), and Decision-making (green) – start with onset of red dot, mask and arrow, respectively. **C**. Illustration of past-present neural interaction calculation. Upper: EEG data at each electrode (left topo) is first regressed to current stimulus and previous report, yielding respective weight maps (*β_current stimulus_*, *β_Previous Report_*). Lower: Each weight map could be denoted as a vector in the 64-dimensional representational space. Past-present interactions are quantified by computing cosine similarity between the vector for current stimulus and the vector for previous report. Positive, zero, and negative values represent aligned, uncorrelated, and flipped relationship, respectively. **D.** Time-resolved grand averaged past-present interactions throughout Encoding (purple), Delay (blue) and Decision-making (green) stages. **E**. Cross-subject correlation between serial dependence behavior (x-axis; regression weight) and past-present neural interaction (y-axis; averaged within significant clusters) during Encoding (left, purple) and Decision-making (right, green) stages. Each dot denotes individual subject. Solid lines denote the best linear fitting. Shaded areas denote 95% confidence interval. For all time-resolved plots, solid lines denote grand average. Shaded areas denote ±1 SEM. Horizontal lines denote significant temporal clusters (cluster-based permutation test, p < 0.05, two-sided, corrected).

First, as shown in Fig. 2B (upper-right panel), information about current-trial target location emerged rapidly during encoding (20-520 ms after stimulus onset) and also appeared during the delay (0-320 ms, 360-490 ms after mask onset) and decision-making stages (60-760 ms after mouse cursor onset; cluster-based permutation test, p < 0.05, two-tailed, corrected). Crucially, previous-trial information (i.e., reported location in the previous trial) could also be decoded from the current-trial neural activities (Fig. 2B, lower-right panel), reactivated during both encoding and decision-making stages (Encoding: 120-260 ms after stimulus onset; Decision-making: 230-300 ms and 330-410 ms after mouse cursor onset). Note that significant clusters in time courses were defined using the same statistical test and criteria unless otherwise specified (cluster-based permutation test, p < 0.05, two-tailed, corrected).

Therefore, previous-trial location information is indeed carried over to the current trial, and the memory reactivations co-occur with current stimulus processing during both the encoding and decision-making stages.

### Two-stage “repulsive-followed-by-attractive” past-present interactions and behavioral correlates

After confirming the co-occurrence of past and present information, we next examined their interactions during both encoding and decision-making stages, focusing particularly on shifting directions, i.e., attractive or repulsive.

Specifically, we examined the neural similarity in representations between past memory reactivation and current stimulus, with positive and negative values denoting attractive (aligned) and repulsive (flipped) bias, respectively (Fig. 2C, lower). To this end, the neural signal of each EEG channel was regressed by three predictors (Current Stimulus, Current Start, Previous report; see Fig. 1B), resulting in a 64-dimensional EEG topographic map of regression coefficients (neural representational axis) for each predictor (Fig. 2C, upper). Next, the cosine similarity between the 64-dimensional representational axis of Previous Report and that of Current Stimulus was computed, signifying the past-present interaction (Fig. 2C, lower). The sign of the past-present interaction value denotes three possible bias directions – aligned (positive value), flipped (negative value), and uncorrelated (near to zero) – corresponding to attractive, repulsive, and non-interactions, respectively (Fig. 2C, lower). This analysis was performed at each time point and on each participant.

As shown in Figure 2D, the past-present interactions exhibited a two-stage dissociated profile: repulsive interactions during encoding (80-370 ms after stimulus onset) and attractive interactions during decision-making (250-290 ms after mouse cursor onset). Furthermore, we examined the behavioral relevance of the two-stage past-present neural interactions, by calculating the Pearson’s correlation coefficients between serial bias behavior (*β_Previous Report_*, see Fig. 1B) and past-present neural interactions (i.e., averaged within significant temporal clusters, see Fig. 2D) across participants, for encoding and decision-making periods, separately. As shown in Figure 2E, only the late attractive interactions correlated with the attractive bias behavior (*r*(28) = 0.37, p = 0.045), but not for the early repulsive interactions (*r*(28) = -0.063, p = 0.75).

Overall, although past-trial information is reactivated during both encoding and decision-making stages, it interacts with present information processing in an opposite two-stage manner: repulsive (flipped) during sensory encoding and attractive (aligned) during decision-making, with the latter further related to attractive serial dependence behavior.

### Past choice instead of past stimulus mediates attractive serial bias

In Experiment 1, stimulus (target location) and report (reproduced location) were highly correlated, making it hard to disentangle the contributions of previous stimulus and previous report to serial bias. Furthermore, it remains unclear how task relevance impacts the observed two-stage process and the underlying brain regions. In Experiment 2, we employed a modified paradigm with MEG recordings to address these questions.

As shown in Figure 3A, participants were presented with two dot stimuli of different colors at two random locations. During decision-making, they recalled the location of one of the two dots based on the retrocue color (Target vs. Non-target dot stimuli), by choosing from two of the response probes closest to the cued target location (Chosen vs. Unchosen response probe) (Fig. 3A, right). Critically, this design could dissociate stimulus and report, since the dot stimuli (Target) and response probes (Chosen) were localized at different locations. Moreover, task-relevance could also be studied based on Chosen vs. Unchosen and Target vs. Nontarget. Twenty-six subjects participated in Experiment 2 and performed well (Fig. 3B; report accuracy: 0.90 ± 0.033; reaction time: 1.77 ± 0.57). Notably, the binary probe choice was converted to a continuous scale (-1 to 1) by incorporating reaction time (RT) as a weighting factor (see Methods for details) to index behavior.

**Figure 3.**
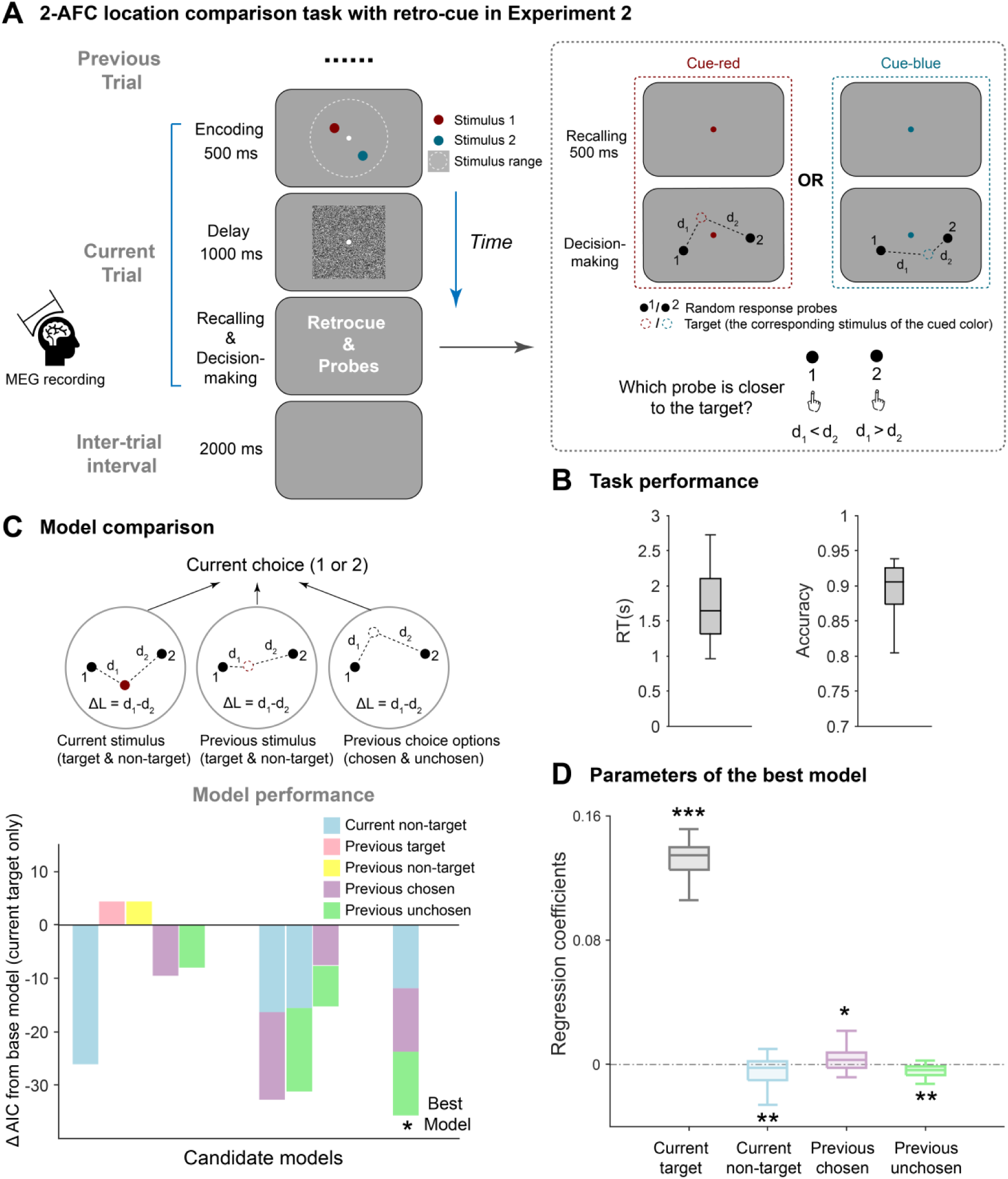
Experimental paradigm and behavioral serial biases (Experiment 2, MEG). **A**. Participants were presented with two colored dots with random location within a 2-D round area, followed by a square mask in the delay period. Afterwards (dotted box in the right), a retro-cue (changing color of central fixation) indicated which dot need be recalled (target vs. non-target). During decision-making, two randomly located response probes (marked by 1 and 2) were presented and participants selected the one closest to the target location. Subjects’ MEG activities were recorded. **B**. Grand averaged accuracy and reaction time (RT). **C**. Model comparison results. Upper: Factors considered to contribute to current-trial choice. Six locations are divided into three categories: current stimulus (target & non-target), previous stimulus (target & non-target) and previous choice option (chosen & unchosen). Lower: ΔAIC of candidate models compared against the base model (current target only). The best model (with the lowest AIC) among all candidates is indicated with an asterisk. Key factors of candidate models are color-coded. **D**. Regression coefficients extracted from the best model. For all the box plots, the bottom edge, central mark and top edge denote the 25th, 50th and 75th percentiles, respectively, and the whiskers extend to the most extreme data points not considered outliers. (*** : p < 0.001, ** : p < 0.01, * : p < 0.05).

First, Linear mixed-effects models (LMMs) with various combinations of factors (Current target, Current non-target, Previous target, Previous non-target, Previous chosen, Previous unchosen) were built to evaluate their contributions to behavior (Fig. 3C). Specifically, candidate models were built in a stepwise manner (gradually adding factors) and compared to the base model (i.e., Current target only), yielding their respective Akaike information criterion (AIC) values with lower value indicating better performance. As shown in Figure 3C (lower panel, star), the model comprising Current target, Current non-target, Previous chosen, and Previous unchosen factors performed best (ΔAIC = -35.57; see Table S1 for model comparison details), suggesting that current choice is biased by past-trial choice (chosen and unchosen) rather than past-trial stimuli (target and non-target). Moreover, regression coefficients extracted from each subject reveal different directions of the four factors biasing the current choice, with positive values (attractive) for Current target and Previous chosen and negative values (repulsive) for Current non-target and Previous unchosen (Fig. 4D; Current target: 0.13 ± 0.016, t(25) = 42.80, *p* < 0.001; Current non-target: -0.0052 ± 0.0092, t(25) = -2.91, *p* = 0.0075; Previous chosen: 0.0039 ± 0.0082, t(25) = 2.43, *p* = 0.023; Previous unchosen: -0.0038 ± 0.0055, t(25) = -3.53, *p* = 0.0016; one-sample *t* test, two-tailed).

**Figure 4.**
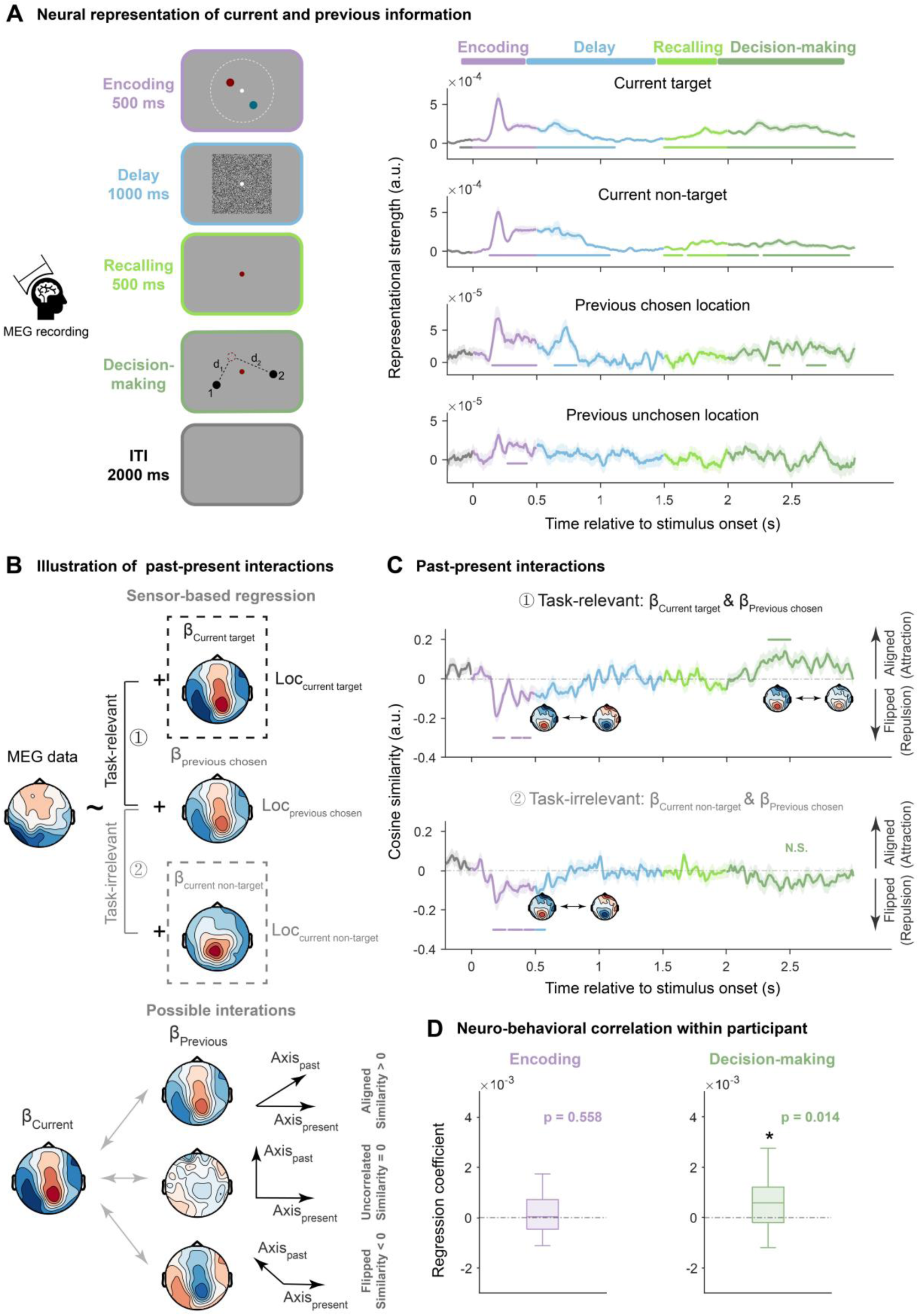
Neural decoding and past-present interactions in Experiment 2 (MEG). **A.** Left: Each trial consists of four stages: Encoding (purple), Delay (blue), recalling (light green), and Decision-making (dark green). Right: Time-resolved grand averaged decoding of current-trial stimulus (target & non-target dot stimuli) and previous-trial choice (chosen & unchosen response probes). **B.** Same past-present neural interaction computation as in Experiment 1 (see Figure 2) was performed in Experiment 2 on MEG sensors. Upper panel: past-present interaction was calculated between weight map of current stimulus (Upper: task-relevant target; Lower: task-irrelevant non-target) and that of previous chosen location (middle), resulting in task-relevant and task-irrelevant past-present neural interactions. Lower panel: three possible interaction directions, with positive, zero, and negative values representing aligned, uncorrelated, and flipped relationship, respectively. **C.** Time-resolved grand averaged past-present neural interaction under task-relevant (between previous choice and current target) and task-irrelevant (between previous choice and current non-target) conditions. The shaded areas denote ±1 SEM. Positive and negative values represent aligned and flipped directions, respectively. **D.** Within-participant correlation between behavioral effect (RT benefit) and past-present neural interaction (averaged within significant clusters) cross trials during encoding (left, purple) and decision-making (right, green) stages, with bottom edge, central mark and top edge denoting the 25th, 50th and 75th percentiles, respectively. The whiskers extend to the most extreme data points not considered outliers. * : p < 0.05. For all time-resolved plots, solid lines denote grand average. Shaded areas denote ±1 SEM. Horizontal lines denote significant temporal clusters (cluster-based permutation test, p < 0.05, two-sided, corrected).

In summary, by employing a modified paradigm in Experiment 2 to separate stimulus and choice information, we demonstrate that the typical attractive serial dependence arises primarily from past-trial choice rather than past-trial stimulus. Moreover, task relevance also modulates serial bias.

### Two-stage reactivations of past choice

We used the same trial-wise RSA decoding method as Experiment 1 to decode variables that essentially contribute to serial dependence (i.e., the best model in Fig. 3C) from MEG activities. As shown in Figure 4A (right), consistent with Experiment 1, target location (Current target) emerged during encoding (0-500 ms after stimulus onset), delay (0-610 ms after mask onset), recalling and decision-making stages (0-500 ms after retro-cue onset; 0-1000 ms after probe onset). Interestingly, although the uncued stimulus (Current non-target) could be discarded after the retrocue, the non-target still emerged during recalling and decision-making (0-140 ms 185-500 ms after retro-cue onset; 0-235 ms 280-1000 ms after probe onset). Thus, current-trial stimulus information is maintained throughout the trial, regardless of task relevance.

In contrast, previous-trial choice reactivations are modulated by task relevancy. Specifically, although the past-trial choice (Previous chosen) was reactivated during encoding (150-500 ms after stimulus onset), delay (140-310 ms after mask onset), and decision-making stages (320-405 ms, 620-765 ms after probe onset), the past-trial unchosen probe (Previous unchosen) was only reactivated during early encoding (270-420 ms after stimulus onset) but not during the late decision-making stage.

Therefore, past-trial reactivation also exhibits a two-stage profile. Both previous chosen and unchosen information is reactivated during early encoding, but only the previous choice is conveyed to the late decision-making stage.

### Task relevance modulates two-stage past-present interactions and their behavioral correlates

Experiment 1 revealed two-stage past-present interaction and its relevance to bias behavior. We used the same approach to calculate neural similarities between past-trial choice and current-trial target/non-target, denoting past-present neural interactions under task-relevant and task-irrelevant conditions, respectively. Similar to Experiment 1, positive and negative values indicate attractive (aligned) and repulsive (flipped) bias (Fig. 4B). As shown in Figure 4C, the early encoding stage (in purple) displayed repulsive past-present interaction for both target (upper; 170-255 ms, 315-385 ms, 405-460 ms) and non-target (lower; 170-265 ms, 290-395 ms, and 415-490 ms after stimulus onset). Meanwhile, during late decision-making (in dark-green), past choice only interacted with current target in an attractive direction (upper; 230-400 ms after probe onset), but not with non-target (lower panel, see Fig. S1 for repulsive interaction between past unchosen and current target during encoding).

We next examined the behavioral relevance of past-present interactions. Motivated by previous studies positing that one benefit of attractive serial bias is to speed up perception and decision-making processes^20^, we used trial-wise reaction time (RT) to quantify serial bias behavior and computed correlations between the inverse of RT (behavioral index) and trial-wise neural interactions (neural index, see details in Methods) within each participant (see Fig. S2 for other results). As shown in Figure 4D, only the late attractive past-present neural interaction correlates with serial bias behavior (*β* = 5.02e − 4 ± 9.62e − 4, t(25) = 2.66, p = 0.014, one-sample t-test, two-tailed), while the early repulsive interaction did not (*β* = 8.38e − 5 ± 7.19e − 4, t(25) = 0.59, p = 0.56, one-sample t-test, two-tailed).

Overall, Experiment 2 confirms the two-stage “repulsive-then-attractive” past-present interactions found in Experiment 1 (Fig. 2D), and further demonstrates the dissociated nature of sensory-encoding and decision-making stages: the early stage is automatic and adaptation-like, and the late stage tends to be modulated by task demands and essentially contributes to typical attractive serial bias behavior.

### Two-stage past-present interactions arise in dissociated brain regions

MEG activities were recorded in Experiment 2 to examine spatiotemporal neural dynamics of past-present interactions. We finally investigated brain regions associated with the two-stage past-present interactions. We first addressed the question at sensor levels (Fig. 5AB), by dividing all the MEG sensors into anterior (green) and posterior (purple) ones, and performed the same analyses on the two halves separately. First, the posterior regions (purple) dominated the early encoding stage (Fig. 5A), exhibiting reactivation of previous choice (150-370 ms, 405-495 ms) as well as repulsive past-present interaction (175-275 ms, 310-390 ms). Meanwhile, the anterior regions (green) drive the late decision-making stage (Fig. 5B), with significant past reactivation (110-410 ms, 605-755 ms) and positive past-present interaction (365-510 ms, 540-635 ms).

**Figure 5.**
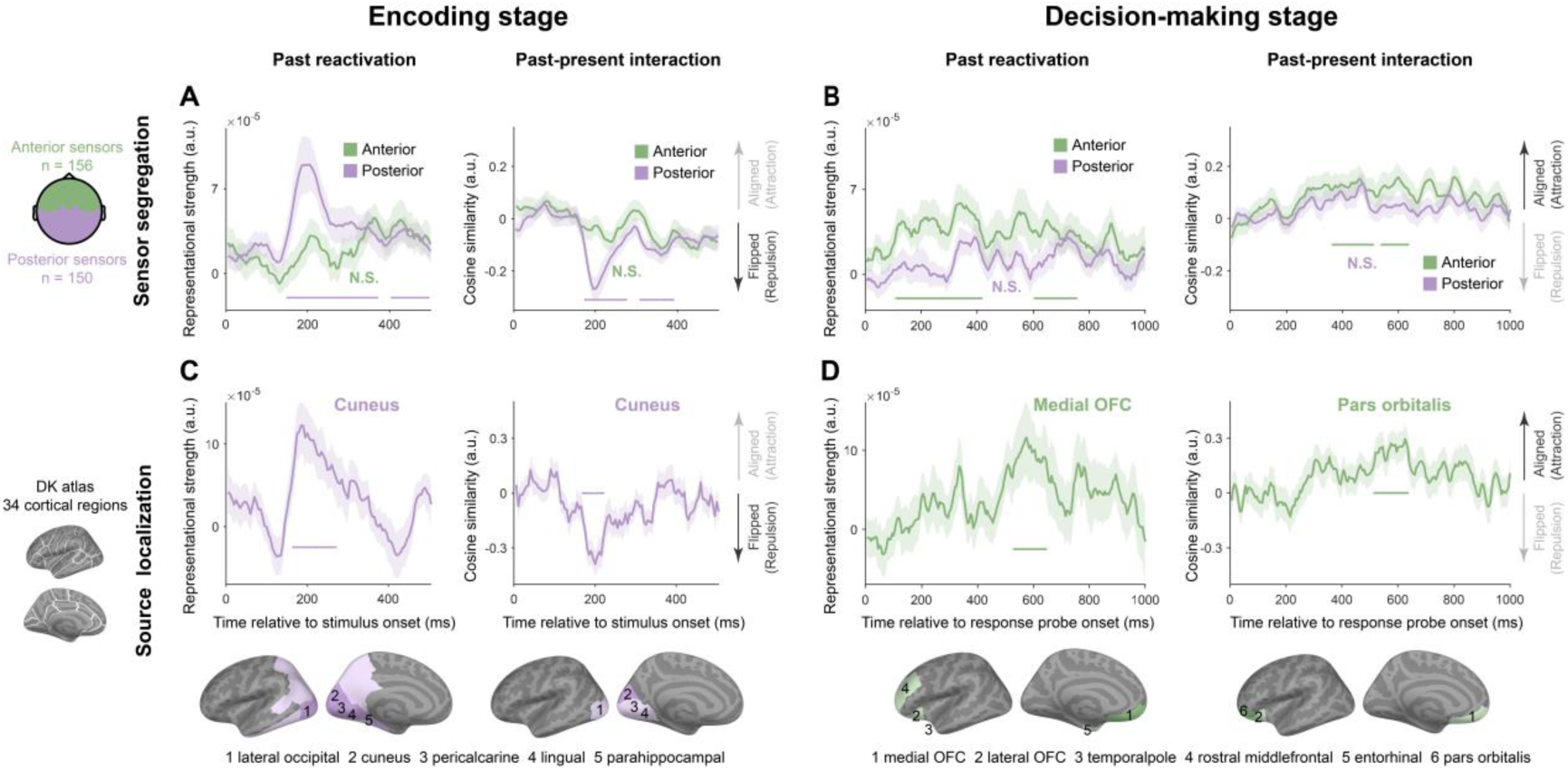
Two-stage cortical origins of serial biases (Experiment 2, MEG). **A-B.** Sensor-based results. 306 MEG sensors are split into 156 anterior (green) and 150 posterior (purple) halves (leftmost, up), within which analyses are performed separately. **A.** Sensor-level results during Encoding stage. Time-resolved grand averaged decoding of previous chosen location information (Left) and past-present neural interactions (Right), for anterior (green) and posterior (purple) halves. Positive and negative values in past-present interaction represent aligned and flipped directions. **B**. The same as A but during Decoding-making state. **C-D.** source-level results. Analyses are performed for each of the 34 cortical regions identified based on the Desikan-Killiany atlas (leftmost, bottom) separately. **C.** Source-level results during Encoding stage. Upper: Time-resolved grand averaged decoding of previous chosen location information (Left) and past-present neural interactions (Right) within cuneus, a representative sensory region. Lower: Significant regions during Encoding stage displayed on inflated cortical surface with their names shown at bottom. Dark and light colors denote temporal clusters with or without spatial corrections. **D**. Same as C but during Decision-making stage. Upper: Time courses within PFC regions (Medial OFC for past reactivation, left; Pars Orbitals for past-present interaction, right). Shaded areas denote ±1 SEM. Color-coded horizontal lines denote corresponding significant clusters (cluster-based permutation test, p < 0.05, two-tailed, corrected). “N.S.” denotes nonsignificant.

We next examined the source level, by performing the same analyses within each of 34 surface regions of Desikan-Killiany Atlas^36^. Consistent with sensor-level results, brain regions mediating past-present interaction during encoding (Fig. 5C, dark purple) were located in posterior areas (cluster-based permutation, p < 0.05, two-tailed, spatial-temporally corrected). Specifically, regions showing significant past reactivations included the lateral occipital cortex, cuneus, peri-calcarine, lingual and parahippocampal areas (Fig. 5C, left, dark blue), and past-present interaction occurred in the cuneus region (Fig. 5C, right, dark blue). In contrast, during late decision-making, anterior brain regions played major roles (Fig. 5D, dark green; cluster-based permutation, p < 0.05, two-tailed, spatio-temporally corrected), with reactivation of previous choice in the medial orbitofrontal cortex (medial OFC; left, dark green), and the past-present interaction in pars orbitalis (right, dark green). Note that light colors denote significant regions without spatial corrections. Importantly, the grand average time course within the corresponding source-level regions displayed temporal profiles commensurate with sensor-level results (Fig. 5CD, upper; see Fig. S3 for time courses in other regions).

Taken together, the two-stage “repulsive-followed-by-attractive” past-present interactions are mediated by dissociated brain regions, i.e., early sensory cortex and prefrontal cortex, respectively.

## Discussion

Serial dependence, as a robust and automatic effect occurring on a wide range of features and paradigms, has been posited to arise from perceptual continuity-field integration^4^ or a two-stage process comprising sensory-level repulsion and post-perceptual integration^11,22^. Here, in two experiments with EEG/MEG recordings, we provide novel neural evidence supporting the latter view. In particular, reactivation of past-trial reports interacts with current information processing in two stages, i.e., first repelling the present during sensory encoding, and then attracting it during decision making, originating from the sensory cortex and prefrontal cortex, respectively. While the early stage occurs automatically, the late stage is modulated by task relevance and predicts serial bias. The two-stage past-present interactions occurring in distinct brain regions might also illustrate the general mechanism of past-present influences in many cognitive functions.

It has been extensively debated at which stage serial dependence occurs^4,5,11,22,23^. Despite large amounts of behavioral studies, neural evidence remains mixed and scarce^25–27,37^. This might be due to the rapid co-evolving information flows across brain regions that are difficult to disambiguate in previous paradigms. Here, by temporally separating encoding and decision-making stages in delayed-response tasks and leveraging the good spatiotemporal resolution of MEG, we demonstrate the neural dynamic of past-present interactions across various cognitive stages and in distinct brain regions, thus unifying previous contradictory findings and advocating the two-stage account of serial dependence.

The early repulsive past-present interaction reflects sensory adaptation that promotes novelty detection based on efficient coding principles^11,22,38^. Recent stimuli, either across-trial or within-trial, are found to repel present sensory processing, especially when stimuli are of high intensity and long duration^4,26^. Visual cortical recordings of rats reveal long-term stimulus-specific adaptations triggered by brief stimuli^39^. Here, the repulsive past-present interactions occur in the sensory cortex, thus consistent with previous findings^26,27,40^. Importantly, we also show that it is past choice (response probe) rather than past stimulus (dot stimulus) that is conveyed to the current trial and engages in the repulsive past-present interactions. The clear dissociation of stimulus and choice arises from Experiment 2 design, whereby stimulus needs to be converted to decision variable during decision-making, so that response probes replace the original stimuli to influence the next trial. Moreover, reactivations of both chosen and unchosen past-trial probes as well as their repulsive interactions with current stimulus in the sensory cortex, which further correlate with repulsive serial bias behavior (Fig. S2), also highlight the automatic nature of low-level efficient coding at the early stage of serial dependence.

The late attractive past-present interaction, in contrast, occurs during decision-making in the prefrontal cortex such as OFC and IFG. This is also in line with previous findings disclosing the involvement of higher-level areas (e.g., PPC and dlPFC) in attractive serial bias^28,29,41–43^. The two-stage model posits that attractive serial bias emerges from post-perceptual Bayesian inference^10,11,22,24,27^, through which prior information is integrated with current information in the prefrontal cortex^44–46^. In addition, task relevance modulates the attractive past-present neural interaction, similar to previous findings^26,47^. Furthermore, the reactivated past choice aligns with target in neural representations but is orthogonal to non-target (nonsignificant past-present interactions). This representational orthogonalization might help minimize interference from task-irrelevant distractors in Bayesian integration^35,48^.

Finally, as past-trial choice is not reactivated by retrocue during working memory (WM) retention but activated during decision-making to mediate attractive serial bias, our results also indicate that the post-perceptual attractive bias occurs during decision-making rather than WM maintenance.

Both repulsive and attractive past-present interactions rely on memory reactivations of past. Consistent with the ‘activity-silent’ WM theory^49^, recent studies show that past-trial information silently stored in WM could be reactivated either before stimulus onset or after^19,28,31–34,50^. Here, only past-trial information that contributes to serial bias behaviors is reactivated, suggesting a substantial role of memory reactivation in serial dependence. Notably, our previous study revealed feature-specific characteristics of past reactivation, i.e., reactivated by corresponding features^19^. Meanwhile, since it is an instantaneous perceptual decision-making task where sensory encoding and decision-making co-evolve, the stages for these feature-specific reactivations could not be identified. Here, by using delayed-response tasks to separate encoding and decision-making stages within a trial, we show clear dissociated stages of past reactivations. Specifically, the failure of retrocue in reactivating past-trial information extends previous findings and suggests that past reactivation is not a fixed pattern but flexible and task-dependent.

Specifically, the retrocue is limited to current stimuli, while the goal-based decision-making stage necessitates activations of past goal-related information (i.e., prior) and present decisional evidence. Finally, past information is retained in sensory and temporal-parietal regions during encoding and frontal-temporal regions during decision-making, supporting distributed neural substrates of WM^51^, and different storage formats for memory, i.e., sensory or abstract representations^52–54^.

To understand the whole loop of serial dependence, in addition to locating its operating stage, identifying the site of prior generation, i.e., where is the information that induces future biases constructed, is also crucial^5^. Converging evidence has shown that prior information causing attractive serial bias is a high-level construct, which is a subjective perception or decision that can incorporate context-information and requires conscious awareness^12,18,19,23,47,55^. However, stimuli and responses are highly correlated, making it challenging to distinguish whether the prior is based on pure report or stimulus. We demonstrated that it is previous reports rather than previous stimulus that contributes to serial bias. Hence, the prior causing attractive bias is a high-level conscious construct derived from task goals.

Taken together, combining EEG/MEG recordings with task paradigms that temporally differentiate cognitive stages and dissociate task variables, we demonstrate the operational circle of serial dependence. The goal-based report formed during decision-making constitutes the prior that biases subsequent perception and decision-making. Sensory traces and abstract codes, stored in different memory-related regions, are reactivated during incoming encoding and decision-making stages, and repulsively and attractively influence current information processing, respectively. This two-stage past-present influence illustrates how the brain balances novelty detection with stable perception over time and provides potential insights into other cognitive functions.

## Methods

### Participants and experimental design

#### Participants

In Experiment 1 (EEG study), we recruited 31 participants, and two were excluded from further analysis because one didn’t finish the task, and one performed poorly. For the remaining 29 participants (13 females), their age ranges from 18 to 27 (mean = 21.6, SD = 1.83). In Experiment 2 (MEG study), 27 participants were recruited and one was excluded from further analysis because of poor performance. The age for the remaining 26 participants (17 females) ranges from 19 to 26 (mean = 21.6, SD = 2.02). All participants in the two experiments have normal or corrected-to-normal visions, and gave written informed consent before commencing the experiments. Both experiments were approved by the Departmental Ethics Committee of Peking University, and were conducted in accordance with the Declaration of Helsinki.

#### Apparatus and materials

Both experimental programs were developed using Matlab (MathWorks Inc., 586 Natick, MA, USA) and Psychophysics Toolbox^56^. In Experiment 1 (EEG study), all visual stimuli were displayed on a 32-inch Display++ LCD screen with a refresh rate of 100 Hz and a resolution of 1920×1080 pixels. Participants sat 57 cm away from the screen and we used a chin-rest to keep their head steady. In Experiment 2 (MEG study), Visual stimuli were presented on a 26-inch rear-projection screen with a resolution of 1024×768 pixels and a refresh rate of 60 Hz. The screen was 75 cm away from the participants. In both experiments, all the visual stimuli were presented on a gray background.

#### Experimental procedure in Experiment 1 (EEG study)

Participants were instructed to reproduce the location of a red dot by moving and clicking a mouse (Fig. 1A). Locations of the red dot and the mouse cursor’s start point were independent, both randomly sampled from a uniform distribution within a 2-D round area (15° visual angle in diameter, edge indicated by the white dashed circle in Fig. 1A). Specifically, in each trial, a red dot (0.4° visual angle in diameter) was first presented for 520 ms, and then masked by a square noise patch (white noise smoothed with a 0.91° standard deviation Gaussian kernel, 15° visual angle in diameter) for 1020 ms, after which a mouse cursor appeared at a random location. Participants had to view and memorize the red dot’s location while maintaining their fixation at a central white dot (0.2° visual angle in diameter) that was presented throughout the trial. After seeing the mouse cursor, they needed to move it to the remembered location of the red dot and left-click. They could adjust their response by moving and clicking it again. When they were satisfied with their response, they could press the ’space’ key to submit their final report and end the trial. There was no time limit for the response. A blank screen with gray background was presented for 2000 ms before the next trial started. Participants first completed a practice session of 10 trials, and then started the formal experiment, which consisted of 32 blocks of 50 trials each, with a one-minute break between blocks. EEG activities were recorded throughout the experiment.

#### Experimental procedure in Experiment 2 (MEG study)

Participants viewed and memorized two differently located and coloured dots (red and blue) and recalled one of them to do a location comparison task (Fig. 3A). In each trial, participants first viewed both red and blue dots, which were independently and randomly drawn from a uniform distribution within a 2-D round area (15° visual angle in diameter, edge indicated using white dashed circle in Fig. 3A), for 500 ms (Encoding), and then encountered a mask of square noise patch (white noise smoothed with a 0.91° standard deviation Gaussian kernel, 15° visual angle in diameter) for 1000 ms (Delay). During the encoding and delay stages, participants had to maintain their fixation on a central white dot (0.2° visual angle in diameter). In the next recalling stage, the color of the central fixation changed to either red or blue, indicating which item had to be extracted from memory and used in the following location comparison task.

We referred to the retro-cued item as “target” and the uncued one as the “non-target”. Two black dots (“probe”, labeled 1 or 2), which were also independently and randomly drawn from a uniform distribution within the same 2D round area as the red and blue dots, were presented on the screen 500 ms after the appearance of the retro-cue, and participants were asked to indicate, by keyboard input (choosing 1 or 2), which probe was closer to the location of the memorized “target”. There was no time limit in their response. A blank screen with gray background was presented for 2000 ms before the next trial started. Each participant practiced 20 trials first, and those with at least 75% accuracy were allowed to enter the formal experiment. The formal experiment consists of 5 blocks of 100 trials each, with one-minute break in between. Throughout the experiment, MEG activities were continuously recorded.

### Behavioral data analysis

#### Multiple linear regression model in Experiment 1 (EEG study)

To examine what factors contribute to participants’ performance (reported location), we fitted a multiple linear regression model to the data (Fig. 1B), which was given by:

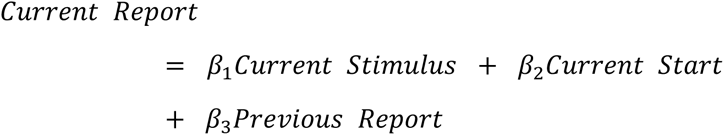

where “Current Report”, “Current Stimulus”, and “Current Start” refer to the reported location, the location of the presented item, and the location of the presented mouse cursor in the current trial, respectively, and “Previous Report” is the reported location in the previous trial. For each participant’s data, the model was fitted for X and Y coordinates separately and the acquired *β*_*x*_ and *β*_*y*_ were averaged to obtain the regression coefficient for each predictor^14^.

The parameter β_3_ reflects the serial bias effect. A positive value indicates an attractive effect, i.e., the reported location in the current trial is biased towards the previously reported location, while a negative value corresponds to a repulsive effect.

#### Linear Mixed Models (LMM) in Experiment 2 (MEG study)

As participants’ report accuracy is high, to quantify their performance in a fine scale, we computed the “weighted choice”, by coding their categorical report, as -1 (probe 1) and 1 (probe 2), and then multiplying it by the rescaled reciprocal of reaction time, which is in the [0,1] interval. Thus, the “weighted choice”, ranges from -1 to 1, with a value closer to the two bounds indicating a more confident choice.

We next evaluated whether the “weighted choice” can be accounted by six types of locations, which belong to three groups, current stimuli (target and non-target), previous stimuli (target and non-target) and previous choice options (chosen and unchosen). As the task is to compare whether the memorized target location is closer to Probe 1 or Probe 2, distances between the target and respective probes, *d*_1_ and *d*_2_, are calculated, and the decision variable, Δ*L*, is the distance difference, *d*_1_ − *d*_2_. Similarly, Δ*L* is computed for each of other locations that possibly affect the “weighted choice”.

Probe locations could bias decisions by themselves. To evaluate this systematic decision bias, for each pair of probes, we randomly sampled a location within the same 2D round area (15° visual angle in diameter) to compute its *ΔL* and repeated this procedure 10000 times to obtain the mean value of *ΔL*(s). The resulting averaged *ΔL* reflects the inherent decision bias by the probes, which is further subtracted from the *ΔL* for each of six candidate locations to obtain the corresponding Δ*L*_*Demean*_.

We fitted linear mixed-effect models to data to examine whether the weighted choice can be predicted by the Δ*L*_*Demean*_ of each candidate location. And a full model is given by:

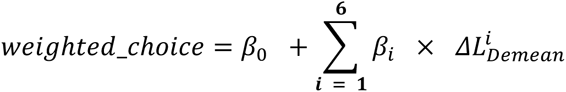

where i refers to each of six candidate locations. The regression coefficient *β*_*i*_ represents the influence of one position on current choice, with a positive value indicating attraction and a negative value indicating repulsion. Model performance was evaluated using Akaike information criterion (AIC) with a smaller value indicating better performance. The base model is the model assuming that the “weighted choice” is only determined by current target location. Models with extra possible combinations of other candidate locations were compared against the base model, and the correspondingΔAIC was calculated. Note, candidate models were built in a stepwise manner by gradually adding factors that make model perform better (lower AIC). The linear mixed-effect models were performed using fitglme function in the Matlab Statistics and Machine Learning Toolbox.

### Neural data analysis

#### Neural data acquisition and pre-processing

We used the 64-electrode actiCAP system and two BrainAmp amplifiers with BrainVision Recorder software (Brain Products) to acquire EEG signals in a sound-attenuated, dimly lit, and shielded room. To record the vertical electrooculography, an electrode was positioned below the right eye. The impedance of all electrodes was kept below 15 kΩ. During the recording, the electrode FCz was used as reference, and all signals were sampled at 500 Hz. The EEG data were pre-processed using FieldTrip toolbox^57^. Data were first epoched into single trials from 0.4 s before stimulus onset to 2.5 s after the stimulus onset. The epoched data were re-referenced to the mean signal of all electrodes and then baseline-corrected with the baseline set from 0.4 s to 0 s prior to the stimulus onset.

Subsequently, EEG data were band-pass filtered within the range of 2 Hz to 45 Hz, and downsampled to 100 Hz. Any channel of excessive noise was replaced with the average value of their neighbouring channels. Finally, to eliminate the eye-movement related and other artifact components from the data, an independent component analysis (ICA) was conducted.

The neuromagnetic signal was acquired using a 306-sensor MEG system (102 magnetometers and 204 planar gradiometers, Elekta Neuromag system, Helsinki, Finland) in a sound-attenuated, dimly lit, and shielded room at Peking University. Both horizontal and vertical electrooculograms (EOGs) were recorded. Before each block, we used online head localization to calibrate the participant’s head with respect to the MEG sensors. Head movements between blocks must remain within 3 mm for further analysis. The spatiotemporal signal space separation (tSSS) technique implanted in MaxFilter was employed to reduce external noise. The MEG signal was sampled at 1000 Hz. For accurate source localization, structural T1-weighted MRI images (1×1×1 mm^3^ resolution) were acquired for each participant on a 3T GE Discovery MR750 MRI scanner (GE Healthcare, Milwaukee, WI, USA). The MEG data were also pre-processed using FieldTrip toolbox^57^. The preprocessing procedure of MEG data is same as what we performed for EEG data except that MEG data were band-pass filtered at 1-40 Hz, and down-sampled to 200 Hz.

### Trial-wise Representational Similarity Analysis (RSA)

We developed a time-resolved multivariate decoding method, i.e., trial-wise Representational Similarity Analysis^58^, to examine neural representations of different types of locations across time. The rationale of this analysis is that neural patterns underlying two locations close to each other are similar and the neural representation similarity decreases as the locations get distant. Linear regression was performed to quantify the relationship between neural representation similarity and location similarity, and the regression coefficient corresponds to the neural representation strength (Fig. 2A). Unlike previous studies employing RSA analysis, which typically have discrete stimulus conditions, locations were randomly sampled within a continuous 2D space in both of our experiments. Thus, we calculated the location and neural similarities for each pair of trials, i.e., a trial-wise analysis. Specifically, we computed the Representational Dissimilarity Matrix (RDM) of location separately for the X and Y axes as follows:

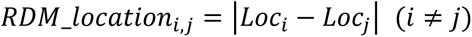

where i and j are trial indices, and Loc represents the X or Y coordinate of a stimulus’ location with the origin being the center of the round area within which stimuli were presented. And at any given time point, the neural RDM is given by:

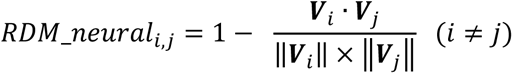

where ***V*** is a vector in the neural representational space at the given time, and i and j are trial indices. In EEG data analysis, the value of each dimension in ***V*** is the EEG signal from a certain channel, and the dimension of ***V*** corresponds to the number of EEG channels. In MEG data analysis, we first performed Principal Component Analysis (PCA) on the 306-dimensional signal and kept the smallest number of principle components that explained 99% variance in total (number of components: 59.1 ± 2.60), and the dimension of ***V*** corresponds to the number of selected PCA components. In practice, we took advantage of spatiotemporal neural dynamics in a predefined time window (50 ms) to increase decoding performance.

Specifically, at each time point, ***V*** is determined by pooling neural signals within 50 ms (centered at the given time point) together as a multivariate pattern. To increase decoding accuracy of previous locations, we also regressed out the influence of current locations on the neural data and used the residual of neural signals. To quantify the neural representation strength, for each type of location, we performed a linear regression:

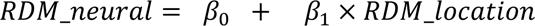

where *β*_0_ is the intercept and *β*_1_ is the regression coefficient for the location. For each participant’s data at each time point, we fitted this model for the X and Y axes separately. The neural representation strength of a certain location at a given time point is obtained by averaging the *β*_*x*_ and *β*_*y*_.

#### Past-present interaction analysis

To evaluate how past information interacts with present neural processing, we examined whether the past reactivation shares similarity with the neural representation of current stimulus. First, for each EEG/MEG sensor, the signal at each time point was regressed onto past and present locations, resulting in n-dimensional vectors of regression coefficients, i.e., neural representation axes, for all predictors (Fig. 2C, Fig. 4B). Specifically, for the EEG experiment, we fitted the following model:

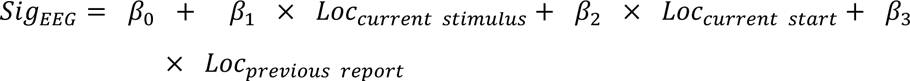

where *Sig_EEG_* is 64-dimensional and Loc denotes the X or Y coordinate for a certain location. The resulting 64-dimensional vectors of *β*_1_, *β*_2_, and *β*_3_ are the representation axes for current stimulus, current start and previous report, respectively. For the MEG experiment, we fitted the following model:

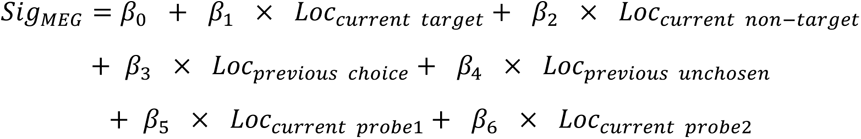

where *Sig_MEG_* is the MEG signal from 204 gradiometers and the resulting six 204-dimensional vectors of regression coefficients indicate the corresponding representation axes.

To quantify the past-present interaction, we calculated the cosine similarity between representation axes of previously reported location and that of the current stimulus location. A negative value indicates that the representational axis of past information is “flipped” in comparison to that of present, corresponding to the neural adaptation-like effect or repulsive serial effect^26^. A positive value indicates that past information is “aligned” with the present, corresponding to the attractive serial effect (Fig. 2C, Fig. 4B).

For each participant’s data at each point, we fitted the regression model separately for the X and Y axes, calculated the respective “past-present interaction,” and then averaged them to acquire the overall interaction. The resulting time series of past-present interaction was then smoothed using a 50 ms sliding time window in both the EEG and MEG experiments.

#### Statistical test for RSA and past-present interaction analyses

For each time period (e.g., encoding) of both experiments, we performed non-parametric sign-permutation test on the time series of “representational strength” and “past-present interaction” to examine whether the variable of interest is significantly different from 0 at the group level^59^. For each permutation, we randomly flipped the sign of the variable being tested from each participant with a 50/50 chance and calculated the group average at each time point. This procedure was repeated 100,000 times to generate a null distribution across time. Next, we compared the original group means against the null distribution at the corresponding time and the p-value was calculated as the percentage of values in the null distribution whose absolute values were larger than the actual mean. We then conducted a cluster-based permutation test for multiple comparisons across time, where a cluster was defined based on a threshold of p = 0.05.

#### Neuro-behavioral correlations

To test whether the past-present interaction is related to behavioral serial bias, we performed neuro-behavioral correlations. Specifically, in the EEG experiment, for each participant, the behavioral serial bias was characterized by the regression coefficient of past report (Fig. 1B), and the neural effect was quantified by averaging the past-present interaction values across significant time clusters during encoding or decision stage. Pearson correlation was performed to examine the neuro-behavioral correlation across participants. In the MEG experiment, we calculated neuro-behavioral correlation within participants (see Fig. S2 for nonsignificant across-participant correlations). The rationale of the analysis is based on the finding that attractive serial dependence could facilitate neural processing and lead to fast response times^20^. The behavioral serial bias was characterized using the reciprocal of reaction time in each trial. For the neural effect, we employed a leave-one-out method to obtain the trial-by-trial past-present interaction. Specifically, for each trial, we fitted a regression model and calculated past-present interactions using all trials except this one. We then derived the trial-level past-present interaction by subtracting the past-present interaction obtained without this trial from the past-present interaction obtained from all trials. The time series of trial-level past-present interactions was smoothed using 50 ms sliding window. Next, we averaged the trial-level past-present interaction values across significant time clusters during the encoding or decision period to get the trial-level neural effect. To quantify the neuro-behavioral relationship, for each participant, we fitted a linear regression model:

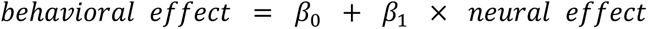

where *β*_1_ characterizes the neuro-behavioral relationship, with a positive value indicating the behavioral benefit of neural serial dependence, i.e., stronger past-present interactions lead to faster response times.

#### Sensor Segregation and source localization analyses of MEG data

To investigate the spatial origins of serial dependence, we first conducted sensor segregation analysis on data from both the encoding and decision-making stages. The MEG sensors were split into two halves and the past chosen location decoding and past-present interaction analyses were carried out on each group of sensors separately to investigate whether there is a spatial dissociation between different effects at different stages. Specifically, we divided sensors based on their Y position (anterior-posterior direction) in the 3-D sensor coordinates. Sensors with positive Y values are classified as anterior sensors (N = 156, 52 magnetometers and 104 gradiometers). Conversely, sensors with negative Y values are classified as posterior sensors (N = 150, 50 magnetometers and 100 gradiometers).

Next, we conducted source localization analysis to directly examine cortical regions implicated in serial dependence. Similar to the sensor segregation, source analyses were also performed on data from the encoding and decision-making stages. For source reconstruction, we first denoised the pre-processed MEG sensor data using PCA and preserved PCA components that explained 95% variance. Subsequently, the MEG data were co-registered with individual’s structural MRI using the MNE_python tool^60^, and Boundary Element Model (BEM) models were generated employing the FreeSurfer watershed algorithm^61^. The forward model was then computed utilizing a surface-based source space with 4096 vertices per hemisphere. Source reconstruction was performed through linear constrained minimum variance (LCMV) beamforming^62^ with a regularization parameter set as 5%, the noise matrix characterized by the baseline data, 0-400 ms before stimulus onset, and data matrices of the encoding and decision-making stages were computed based on the corresponding stage’s data. Then, MEG sources were anatomically mapped to 34 regions of interest encompassing both left and right hemispheres based on the Desikan-Killiany atlas^36^. We combined left and right hemispheres together and conducted the decoding and past-present interaction analyses on each region, just as what we did in sensor analyses but treated a voxel as a sensor here. To address multi-comparison issue across cortical regions, temporal clusters in each region were compared against permuted clusters across all regions, resulting in a spatially corrected p value.

## Acknowledgments

This work was supported by the STI2030-Major Project 2021ZD0204100 (2021ZD0204103 to H. L.), National Natural Science Foundation of China (31930052 to H.L), and China Postdoctoral Science Foundation (2020M680166 to H.Z.). H.Z was supported in part by the Postdoctoral Fellowship of Peking-Tsinghua Center for Life Sciences.

