## supplemental information for "Reactivated past decisions repel early sensory processing and attract late decision-making"

### Supplementary information

#### Table S1

Behavioral model comparison of Experiment 2

| No. (Left to right) | Variables (√ : included) | | | | | | *AIC* | △ $AIC$ from baseline model | Notes |
|---|---|---|---|---|---|---|---|---|---|
| | Current target | Current non-target | Previous target | Previous non-target | Previous chosen | Previous unchosen | | | |
| baseline | √ | | | | | | 20350 | 0 | |
| 1 | √ | √ | | | | | 20323 | -26.09 | |
| 2 | √ | | √ | | | | 20354 | 4.39 | |
| 3 | √ | | | √ | | | 20354 | 4.37 | |
| 4 | √ | | | | √ | | 20340 | -9.61 | |
| 5 | √ | | | | | √ | 20341 | -8.06 | |
| 6 | √ | √ | | | √ | | 20317 | -32.77 | |
| 7 | √ | √ | | | | √ | 20318 | -31.17 | |
| 8 | √ | | | | √ | √ | 20334 | -15.22 | |
| 9 | √ | √ | | | √ | √ | 20314 | -35.57 | Best Model |

**Figure S1**

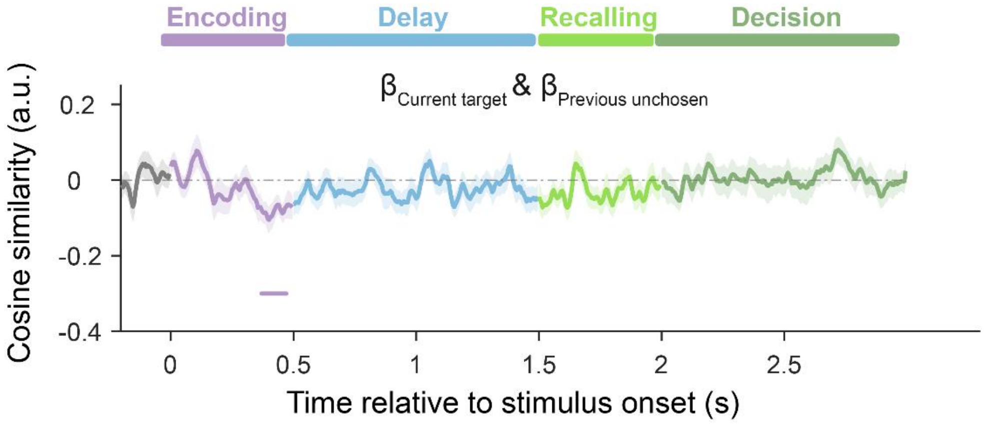


**Figure S1. Time-resolved grand average of past-present interactions between current target and previous unchosen location in Experiment 2 (MEG)**. The shaded areas correspond to ±1 SEM. Positive and negative values correspond to aligned and flipped direction, respectively. Horizontal lines denote significant temporal clusters (cluster-based permutation test, $p < 0.05$, two-sided, corrected).

#### Figure S2

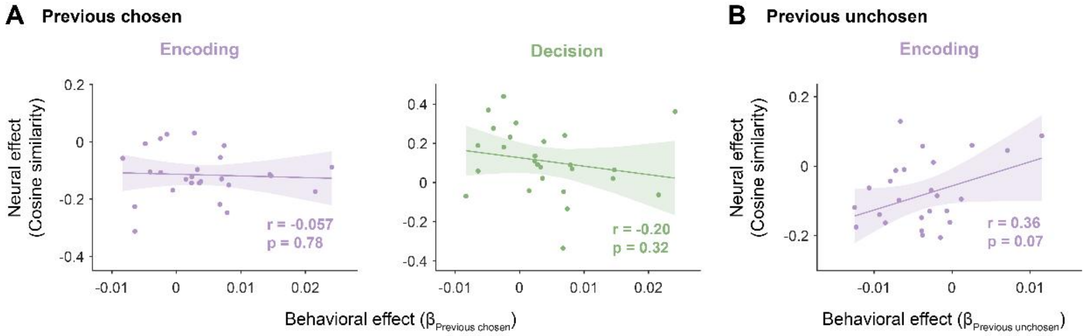


**Figure S2**. **Cross-subject neuro-behavioral correlation in Experiment 2 (MEG). A**. Cross-subject correlation between serial dependence behavior (x-axis; regression coefficient) and past-present neural interaction (y-axis; averaged within significant clusters) induced by previous chosen location during Encoding (purple) and Decision-making (blue) stages. **B**. The same as A but for serial dependence behavior and past-present neural interaction induced by previous unchosen location during Encoding stage. Each dot represents individual participant. The solid lines correspond to the best linear fitting, and the shaded areas correspond to 95% confidence interval.

#### Figure S3

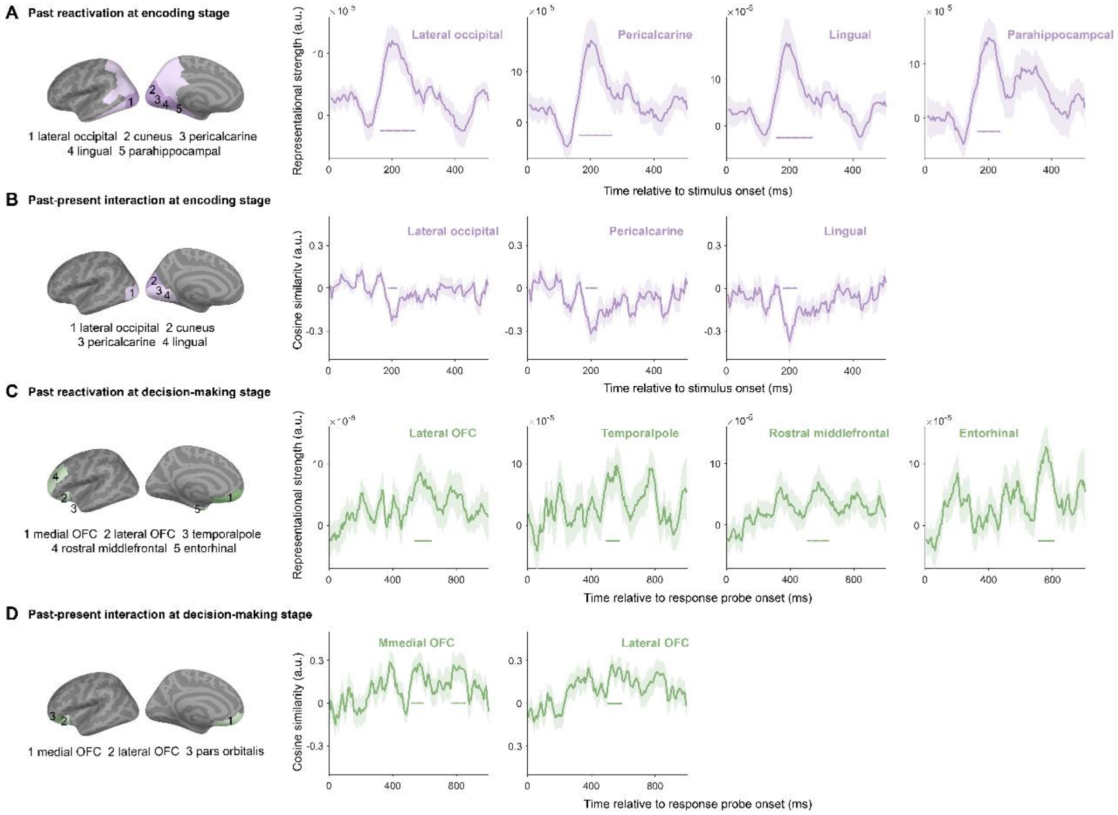


**Figure S3**. **Time courses of two-stage cortical origins of serial biases in all significant regions in Experiment 2 (MEG). A.** Time-resolved grand averaged decoding results of previous chosen location during encoding stage. The regions of significant temporal clusters displayed on the inflated cortical surface and their names are shown at the leftmost. Dark and light colors denote temporal clusters with or without spatial correction. Except Cuneus (shown in Figure 5), time courses of all regions with significant temporal clusters are plotted. **B.** Same as A but for the past-present interactions. **CD.** Same as AB but during decision-making stages. Panel C does not include the result of medial OFC and panel D does not include pars orbitalis. The shaded areas correspond to ±1 SEM. Color-coded horizontal lines denote significant temporal clusters (cluster-based permutation test, $p < 0.05$, two-tailed, corrected).
